# NTM-DB: A Comprehensive Non-Tuberculosis Mycobacteria Genomic Database

**DOI:** 10.1101/2025.07.22.666127

**Authors:** Tianyi Lu, Cuidan Li, Haobin Wei, Yadong Zhang, Zhuojing Fan, Xiaoyuan Jiang, Jie Wang, Peihan Wang, Kang Shang, Yuting Huang, Hongwei Yang, Bahetibieke Tuohetaerbaike, Ying Li, Haitao Niu, Wenbao Zhang, Hao Wen, Yongjie Sheng, Jingfa Xiao, Fei Chen

**Affiliations:** National Genomics Data Center, China National Center for Bioinformation, Beijing 100101, China; Beijing Institute of Genomics, Chinese Academy of Sciences, Beijing 100101, China; University of Chinese Academy of Sciences, Beijing 100049, China; State Key Laboratory of Elemento-Organic Chemistry, College of Chemistry, Nankai University, Tianjin 300071, China; State Key Laboratory of Pathogenesis, Prevention and Treatment of High Incidence Diseases in Central Asia, Clinical Medicine Institute, The First Affiliated Hospital of Xinjiang Medical University, Urumqi 475000, Xinjiang, China; Key Laboratory of Viral Pathogenesis & Infection Prevention and Control (Jinan University), Ministry of Education, Guangzhou 510632, China; Key Laboratory for Molecular Enzymology and Engineering of Ministry of Education, School of Life Sciences, Jilin University, Changchun 130012, China

**Keywords:** Non-Tuberculous Mycobacteria, Genomics, Database, Integrative analysis, Interactive Web Platform

## Abstract

Non-tuberculous mycobacteria (NTM) are a major group of environmental bacteria, with approximately one-third of them causing serious human infections, particularly respiratory diseases. The global rise in the prevalence and severity of NTM infections has posed a major public health challenge. While high-throughput sequencing has generated vast genomic data on NTM, there remains a lack of comprehensive resources for cross-species genomic analysis. To address these limitations, we have developed a specialized database called NTM-DB (https://ngdc.cncb.ac.cn/ntmdb) tailored for NTM researchers and clinicians. NTM-DB offers the most comprehensive collection of NTM genomic and bioinformatic resources, including 16,469 genome assemblies (13,134 newly assembled genomes), 189 type/standard strain genomes representing 177 species and 12 subspecies, 705 MLST types, 33,240 resistance genes, and 74,315 virulence genes. A user-friendly interactive website has been constructed to enable efficient browsing, MLST profiling, searching, online analysis, and downloading of the aforementioned data. Notably, with online analysis tools, users can perform customized genotyping, cross-species phylogeny, pan-genome, and virulence and drug resistance gene annotation analyses using our data and/or their uploaded data. Overall, with its comprehensive data, intuitive interface, and powerful analysis tools, NTM-DB serves as an important resource and reference for NTM researchers and clinicians, improving diagnosis and treatment for various NTM-related diseases, and supporting both scientific discovery and clinical practice.

## Introduction

Non-tuberculous mycobacteria (NTM) are a major category of environmental bacteria [1, 2], with approximately one-third capable of infecting humans, primarily targeting lung tissue and causing various respiratory diseases [3, 4]. Additionally, in the past decade, there has been a notable increase in reported cases of extrapulmonary NTM infections, causing serious conditions such as keratitis, endocarditis, meningitis, sepsis, and even death in severe cases [5]. Notably, there has been a significant global increase in NTM infections in recent years [6], along with the emergence of new pathogenic NTM species, which pose a substantial public health challenge [7, 8]. For example, recent studies from Europe and America have indicated a roughly 25% rise in NTM infections often linked to environmental changes [9–12]. Similarly, major Chinese cities like Beijing, Shanghai, Shenzhen, and Hangzhou have seen NTM detection rates rise by about 30% due to urbanization [13].

The pathogenesis of NTM is complex and varies significantly among different NTM species and molecular types [14]. This diversity leads to a wide range of symptoms that often mimic other illnesses, making accurate diagnosis challenging [15, 16]. For example, currently, about 30% of NTM infections are initially mistaken for multidrug-resistant tuberculosis due to the limitations of routine diagnostic methods such as physiological and biochemical markers and imaging techniques [17].

The rapid development of high-throughput sequencing technology has provided excellent tools and opportunities for the precise diagnosis and treatment of various types of NTM [18]. This has also led to which led to extensive genomic studies of NTM and related infections, generating a significant amount of genomic data [19–21]. To fully utilize this wealth of data, it is urgently needed to integrate cross-species NTM genomic and bioinformatic data (including genotyping, evolution, drug resistance, and virulence), and develop a comprehensive online analysis platform, which will provide a deeper understanding of NTM pathogenicity and offer essential support for researchers and healthcare professionals navigating the complexities of NTM infections.

To date, although there are several microbial databases (such as BV-BRC, [MicroScope](https://mage.genoscope.cns.fr/microscope/home/index.php), [pubMLST](https://pubmlst.org/), [BacWGST](http://bacdb.cn/BacWGSTdb/), and [GTDB](https://gtdb.ecogenomic.org/)) that include some NTM genomic and bioinformatic data, no specialized NTM database has been reported. These databases only collect limited NTM strains/species (fewer than 3000 strains and 100 species) and neglect numerous NTM species/strains, due to their reliance on assembled NTM genomes from GenBank/RefSeq. Therefore, a substantial amount of raw sequencing data from related studies is excluded, resulting in a limited representation of NTM. Additionally, since this NTM information is mixed with immense amounts of data from other microorganisms, it brings inconvenience to the researchers specifically studying NTM. On the other hand, the abovementioned databases have their own focuses. BV-BRC and [MicroScope](https://mage.genoscope.cns.fr/microscope/home/index.php) only provide genomic annotations at the individual strain level, while [pubMLST](https://pubmlst.org/), [BacWGST](http://bacdb.cn/BacWGSTdb/), and [GTDB](https://gtdb.ecogenomic.org/) primarily focus on genotyping. Overall, the integration of cross-species NTM genomic/bioinformatic data (such as genotyping, evolution, drug resistance, and virulence), and related online analysis platforms are still lacking in the NTM research field.

To address the limitations, we have developed a specialized database called NTM-DB (https://ngdc.cncb.ac.cn/ntmdb) tailored for NTM researchers and clinicians. The primary objective of NTM-DB is to encompass the most comprehensive collection of NTM genomic and bioinformatic data **(Figure 1)**. Specifically, NTM-DB includes a total of 13,134 newly assembled whole-genomes and 3,335 GenBank/RefSeq assemblies, covering 177 out of 185 NTM species. Notably, NTM-DB collected the most comprehensive NTM reference genomes for genotyping, including 189 type/standard strain genomes (177 species and 12 subspecies) and 177 representative genomes with optimized high-quality assemblies for each species. It also includes 7,152 records of associated antibiotic susceptibility test results. Secondly, NTM-DB offers customized analyses of multi-locus sequence typing (MLST), cross-species phylogeny, pan-genome, and virulence and drug resistance gene annotation using our data. Correspondingly, it also provides a range of visualization and interactive functionalities by integrating related bioinformatics resources. Additionally, with the use of online analyzing tools, users can perform genome-typing and evolutionary analysis based on our type strains, drug resistance, and virulence gene annotations with their own genomic data. Overall, NTM-DB stands as a crucial resource for advancing NTM research, assisting the clinical diagnosis and treatment of various NTM-related diseases, and contributing to the advancement of scientific discovery and applied innovation within the field.

**Figure 1.**
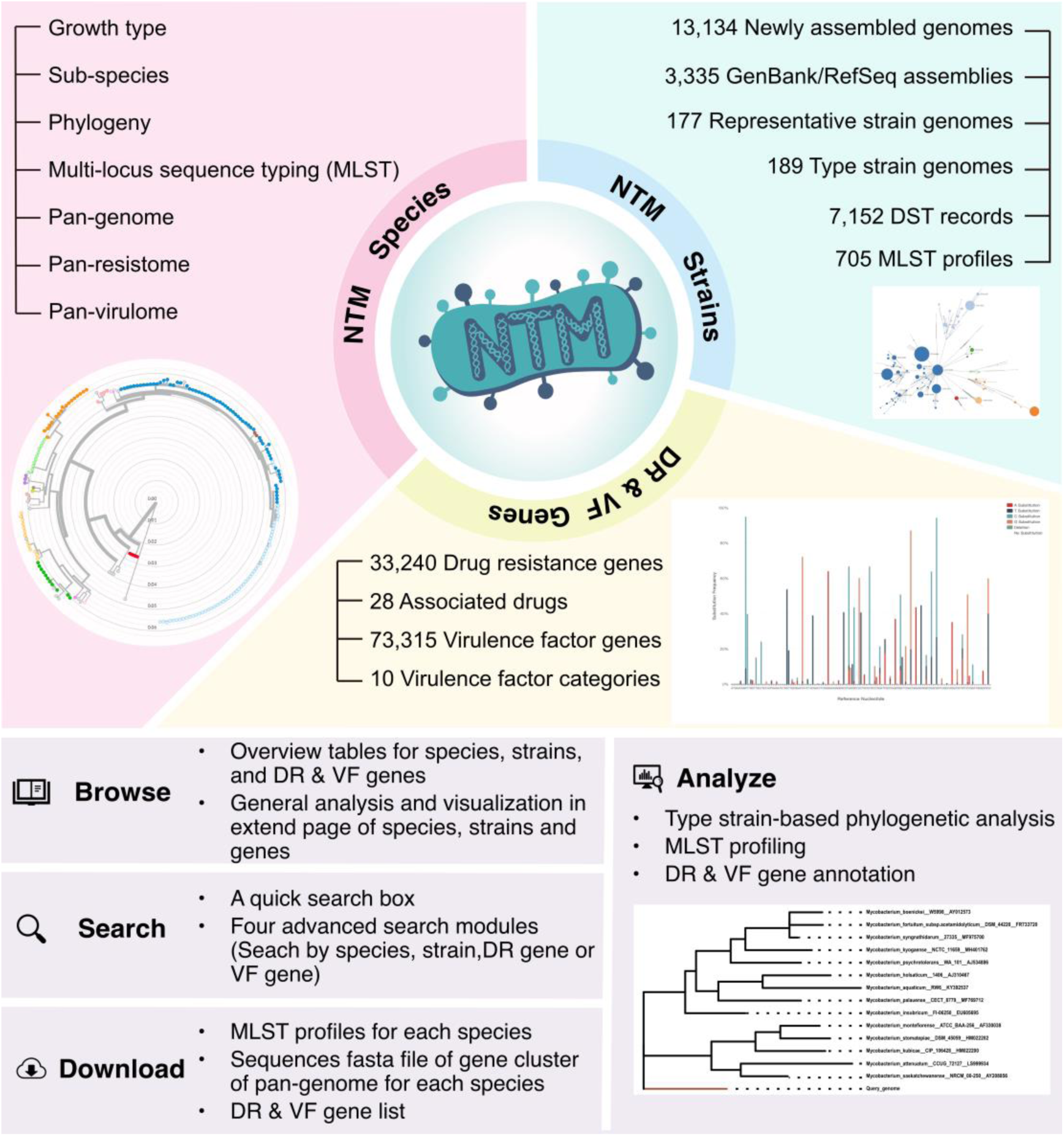
Overview of NTM-DB Contents and Functional Modules. NTM-DB integrates data content and analytical tools focusing on three core objects: species, strains, and genes. The interface is structured into four main functional modules: **(1) Browse**, offering overview tables for species, strains, drug resistance genes, and virulence genes, along with comprehensive visualizations on extended pages; **(2) Search**, featuring a quick search box and four advanced search modules allowing targeted searches by species, strains, drug resistance genes, or virulence factor gene; **(3) Download**, where users can access multi-locus sequence typing (MLST) profiles for each species, download sequences in FASTA format for each gene cluster, and obtain lists of drug resistance and virulence factor genes; **(4) Analyze,** providing tools for type strain-based phylogenetic analysis, MLST profiling, and annotations of drug resistance genes and virulence factor genes. This framework is designed to support dynamic and intuitive interaction with extensive NTM datasets, facilitating detailed genetic and epidemiological studies.

## Database Construction

### Data collection

The NTM-DB database (https://ngdc.cncb.ac.cn/ntmdb) was curated to encompass the most comprehensive collection of NTM genomic and bioinformatic data to date. First, we identified 14,116 strains by reviewing the genomic literature on NTM. During this process, we also collected drug susceptibility test results for 598 strains across 27 drugs, totalling 7,152 data results. Additionally, we retrieved 5,773 pre-publication submission records from the SRA database. For both sets of records, we performed de novo genome assembly and then filtered them based on genome assembly quality and a > 85% average nucleotide identity (ANI) comparison with type strains of NTM species, ultimately resulting in 13,134 newly assembled high-quality whole-genome assemblies. Secondly, we downloaded 3,335 assembled strain genomes from GenBank [22]/RefSeq [23, 24]. In total, 16,469 assembled strain genomes were incorporated into our database, encompassing 177 out of the 185 species officially named by the ICNP [25]. Corresponding metadata (such as associated diseases, geographic origins, and hosts) for the strains was also integrated into the database.

### Data processing workflow

The NTM-DB analysis workflow **(Figure 2)** consists of two main parts: strain-level annotation and species-level integration and comparison. At the strain level, genome sequences are processed through *de novo* assembly and variant calling, followed by annotation for identifying drug resistance and virulence genes. At the species level, we construct phylogenetic trees using the Nextstrain [26], analyze pan-genomes, and identify gene clusters related to drug resistance and virulence factors. This integrated approach enables multi-dimensional data analyses, focusing on evolution, MLST typing, resistance, and virulence.

**Figure 2.**
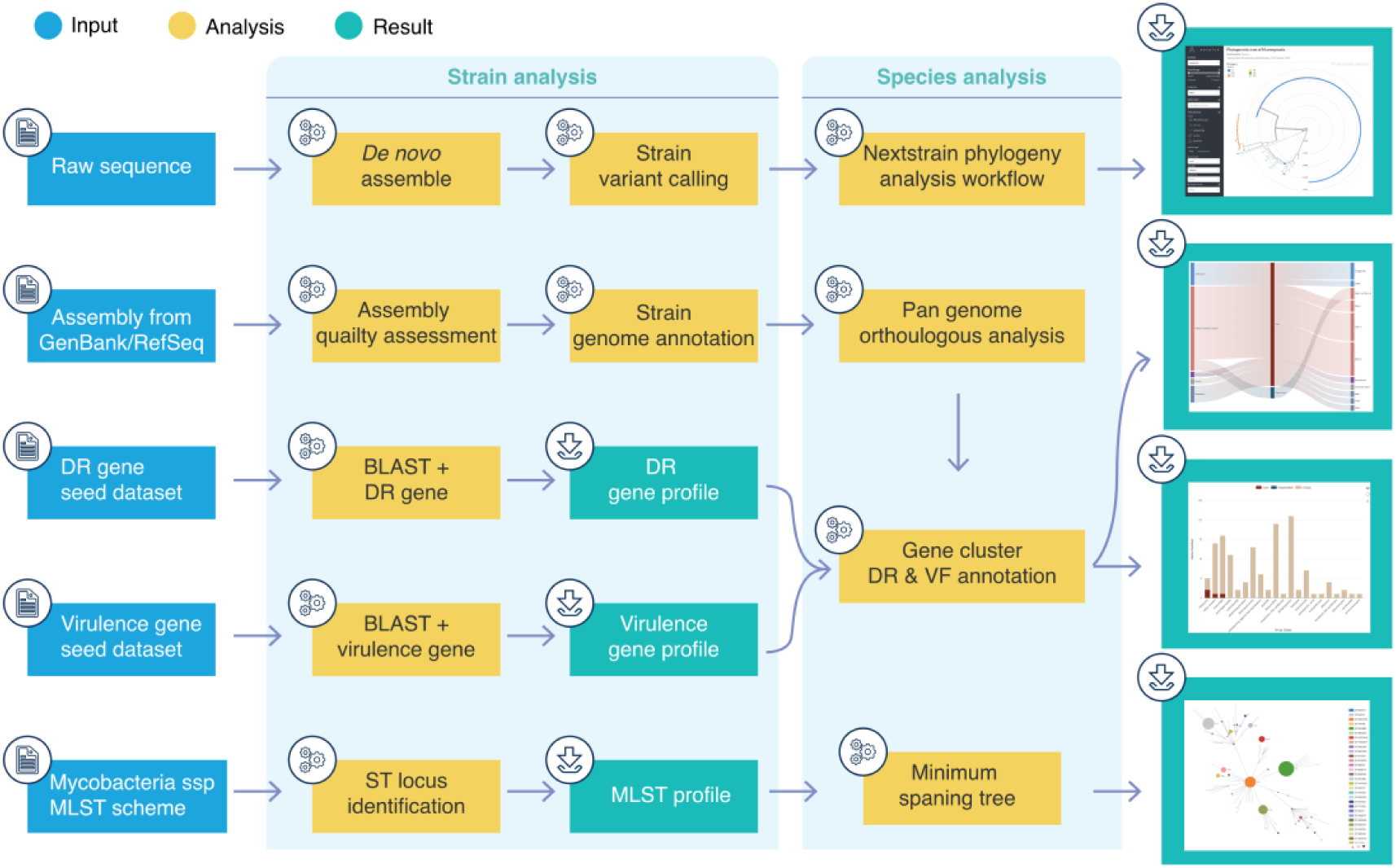
Data Processing Workflow of NTM-DB. The workflow is organized into two main sections, highlighted with light a blue background. The left section represents the strain-level analysis, which includes *de novo* assembly, variant calling, and annotation of drug resistance and virulence genes. The right section illustrates the species-level analysis, encompassing phylogeny construction, pan-genome analysis, and identification of gene clusters associated with drug resistance and virulence factors.

### Genomic data processing

To characterize the bacterial genomes, we integrated two sets of genomic data: one part consisted of raw sequence reads of second-generation sequencing(NGS) samples sourced from the SRA [27, 28], DDBJ [29], and EBI [30] databases, and the other part comprised assembled genomes retrieved from GenBank [22]/RefSeq [24]. For raw sequence files from the first dataset, initial quality control was performed using the fastp tool [31] with the following parameters: qualified quality phred score ≥15, unqualified base percentage limit ≤40%, N base limit ≤5, and minimum read length ≥15 bp. Adapter trimming was performed automatically for both single-end and paired-end data through fastp’s built-in adapter detection algorithm. The pre-processed reads were then assembled *de novo* using SPAdes [32] with optimized parameters including k-mer sizes of 23, 35, and 55, with the --careful flag enabled for error correction. Strains that failed assembly were excluded from further analysis. The assembly quality was subsequently assessed with the QUAST tool [33], applying stringent quality control thresholds: N50 ≥40,000 bp, and L50 ≤60 contigs to ensure that only high-quality assemblies with appropriate contiguity and expected genome size ranges for NTM species were included. After obtaining assembled genomes from both parts, we assessed the completeness and contamination levels of these assemblies using CheckM [34]. Following the completion of the assembly and an extensive quality control regimen for each strain, the assembled genomes were subjected to a series of analyses, including phylogenetic analysis, MLST typing, pan-genomic analysis, and annotation of antibiotic resistance and virulence genes.

### Phylogenetic analysis of species

To delineate the evolutionary relationships within each species and to identify sub-species classifications, we constructed phylogenetic trees based on whole-genome sequencing data for species with more than five strains.

For species among them with reference genomes assembled at the complete level, the Nextstrain Augur framework [26] was used to perform phylogenetic analyses. Specifically, all the strains were mapped to their reference genomes, and single nucleotide polymorphisms (SNPs) were identified by MUMmer 4.0 [35] based on the whole genome sequences. Subsequently, the Maximum Likelihood phylogenetic trees were constructed using IQ-TREE [36]. The phylogenetic tree was visualized using the Nextstrain Auspice platform, providing an interactive interface for exploring phylogenetic relationships. The platform facilitates user engagement with various evolutionary tree visualizations, such as options to colour-code and filtering nodes according to strain metadata.

Additionally, for the 44 species with reference genomes only assembled at the contig or scaffold levels, which are incompatible with Nextstrain’s Augur framework and Auspice visualization, phylogenetic analyses were conducted based on core genes. Here, core genes were identified using Roary [37], followed by a multiple sequence alignment. The phylogenetic trees were constructed using IQ-TREE, and visualized by ggtree [38].

### MLST typing analysis

Multi-locus sequence typing (MLST) analysis was performed (https://github.com/tseemann/mlst) based on a scheme from the pubMLST [39] database. A minimum spanning tree (MST) was subsequently constructed to delineate the genomic relationships among the STs and strains using GrapeTree [40]. To further improve the dynamism and interactivity of the visualization, we migrated the GrapeTree visualization to an Echarts graph.

### Pan-genomic analysis and annotation of virulence and antimicrobial resistance genes

To comprehensively elucidate the functional gene features of each species, a pan-genome analysis was performed for each species. Initially, all the genomes of a specific species were annotated using Prokka v1.14.6 [41]. Subsequently, pan-genome analysis of orthologous gene clusters was conducted utilizing Roary v3.13 [37]. Gene clustering results were used to construct gene family matrices, classifying these gene clusters into three groups based on their pan-genome conservation levels: 1) core gene clusters, consisting of homologous genes present across all analyzed strains of a given species; 2) unique gene clusters, containing genes specific to individual analyzed strains; and 3) dispensable gene clusters, consisting of genes present at least two strains of a given species.

### Annotation of antimicrobial resistance and virulence genes

To characterize the antibiotic resistance and virulence traits of the strains, five databases were utilized: NCBI AMRFinderPlus [42], CARD [43], ResFinder [44], MEGARes [45], and VFDB [46]. Seed datasets for antibiotic resistance genes and virulence genes were constructed, and target sequences were searched against these datasets using BLAST+ v2.12.0 [47]. Subsequently, gene cluster matrices were constructed based on the presence or absence of the annotated antibiotic resistance and virulence genes, categorizing them into core, unique, and dispensable gene clusters according to their conservation levels. The results of pan-resistance and pan-virulence genes were visualized using bar charts and Sankey diagrams, respectively.

### Database implementation

The NTM-DB was constructed using the Django framework (https://www.djangoproject.com/) and deployed via the Gunicorn WSGI server (https://gunicorn.org/). Multiple technologies were incorporated for the front end, including AJAX (Asynchronous JavaScript and XML), Bootstrap (https://getbootstrap.com), CSS (Cascading Style Sheets), Semantic UI (https://semantic-ui.com), Select2 (https://select2.org/), JQuery (https://jquery.com), HTML (HyperText Markup Language), and AG-Grid (https://www.ag-grid.com/). ECharts was utilized for data rendering and visualization, while MySQL served as the database engine for data storage and querying.

## Database Content

### General characteristics of NTM species

Our NTM-DB database includes 16,469 NTM whole-genome assemblies, covering 177 species, with the majority being human-derived (11,033 strains), followed by animal-derived (1,101 strains). *Mycobacterium (M.) abscessus* is the most prevalent species, represented by a substantial collection of 9,797 strains **(Figure 3A)**. This is followed by *M. avium* and *M. intracellulare*, with 2,271 and 1,620 strains, respectively. These results are in agreement with clinical observation that identifies *M. abscessus*, *M. avium*, and *M. intracellulare* as the predominate NTM species causes of pulmonary infections [48, 49]. *M. abscessus* and *M. intracellulare* demonstrate a preference for human infection, with 83% and 70% of strains originating from human sources, respectively **(Figure 3B)**. In contrast, despite the close genetic relationship between *M. avium* and *M. intracellulare, M. avium* exhibits a divergent pattern, with 45.6% of its strains originating from animal sources and 39.8% from human sources. *M. ulcerans* ranking fourth (511 strains), is known for causing skin ulcerative infections through contact rather than airborne transmission [50, 51]. The database also includes other main clinically significant species such as *M. smegmatis*, *M. chelonae*, *M. kansasii*, and *M. marinum*. Collectively, these top nine NTM species account for 90% of all sequenced NTM strains in the database.

**Figure 3.**
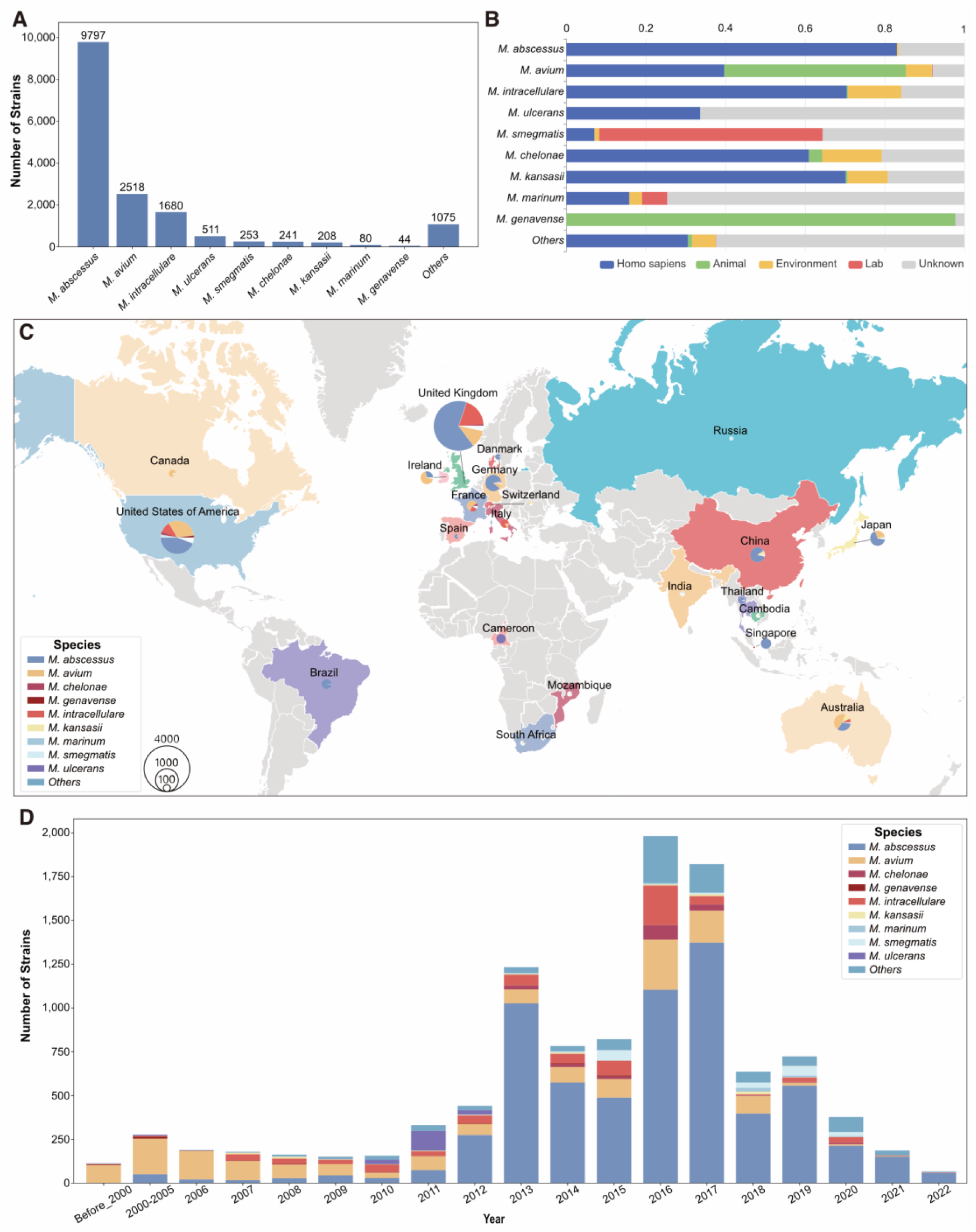
General Characteristics of Major NTM Species in the NTM-DB. A. Species composition of NTM strains. **B.** Host distribution of the top nine NTM species. **C.** Global distribution of the main NTM species. **D.** Temporal distribution of the main NTM species.

### Geographic and temporal distribution of NTM species

Our dataset encompasses 42 countries across five continents, with 21 countries reporting exceeding 50 NTM strains each **(Figure 3C)**. The United Kingdom (4,763 strains), the United States (2,086 strains), Australia (552 strains), Germany (502 strains), and Japan (434 strains) emerge as the top five countries with the highest strain count. These nations exhibit high proportions of *Mycobacterium (M.) abscessus*, with percentages reaching 65.4% in the UK, 44.2% in the US, 32.4% in Australia, 85.7% in Germany, and 66.4% in Japan.

Regarding temporal distribution **(Figure 3D)**, our strain collection spans from 1901 to 2023, with the majority of strains collected after the year 2000 (98.9%) and a peak of 1,981 strains in 2016. Before 2010, *M. avium* was the most prevalent species (60.8%), while *M. abscessus* accounted for only 15.6%. However, *M. abscessus* exhibited rapid growth post-2011, escalating from 22.35% in 2011 to 86.6% in 2022. The significant prevalence of *M. abscessus* in these countries, coupled with its rapid expansion after 2010, aligns with several localized outbreaks worldwide of dominant circulating clones (DCCs) [52, 53], which have evolved increased pathogenicity and drug resistance [21, 54], particularly noted in the UK [19, 55] and the USA [56–58].

### MLST characteristics of NTM species

The MLST (Multi-Locus Sequence Typing) analysis **(Figure 4A, 4B)** identifies the top three sequence types (STs), namely ST5 (2,486 strains, 17.3%), ST3 (1,458 strains, 10.1%), and ST33 (1,276 strains, 8.9%), all associated with *M. abscessus*. These are followed by ST8 and ST81, linked to *M. avium* (1,020 strains, 7.1%) and *M. intracellulare* (1,019 strains, 7.1%), respectively. Notably, *M. abscessus* and *M. chelonae* exhibit a higher diversity of ST types and a lower proportion of dominant ST types, indicating greater genetic variability within these species. Conversely, four major species (*M. intracellulare*, *M. ulcerans*, *M. kansasii*, and *M. genavense*) are dominated by a single ST type, accounting for over 70% of strains, suggesting more conservative genetic characteristics and slower evolutionary rates.

**Figure 4.**
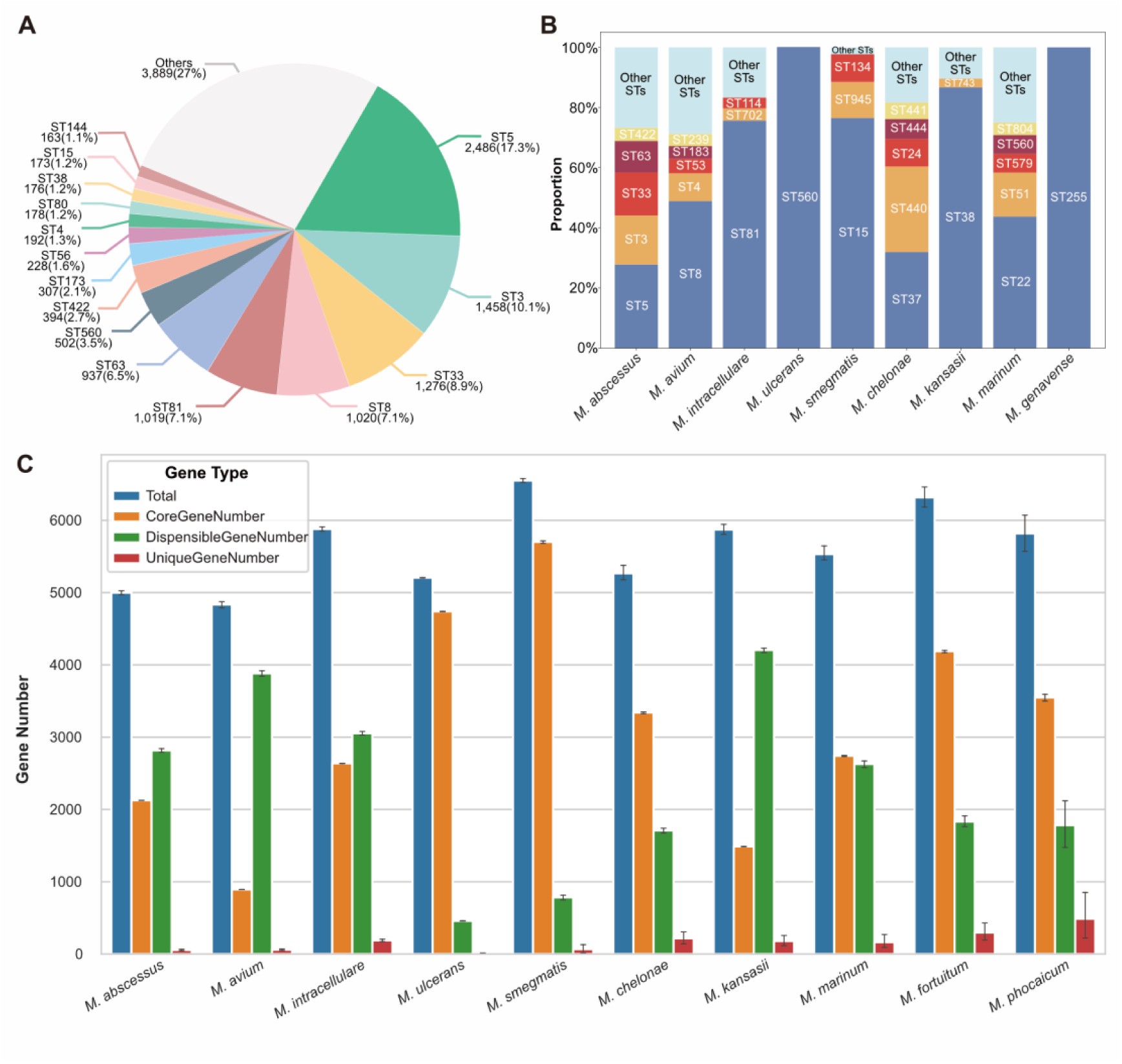
MLST and Pan-Genome Analyses of NTM-DB Strains. A. Distribution of primary STs in the NTM-DB, illustrating the prevalence of various STs. **B.** Composition of STs within the top nine NTM species, highlighting the diversity and distribution across different species. **C.** Mean gene numbers per strain in the top ten species, categorized by Total, Core, Dispensable, and Unique genes based on Pan-Genome analysis. Error bars indicate confidence intervals.

### Pan-genome characteristics of NTM species

Our database includes a pan-genome analysis of 62 species, revealing an open pan-genome for all NTM species. The pan-genome characteristics of the top ten species are shown in **(Figure 4C**). Notably, *M. ulcerans* has the highest proportion of core genes, with 4,739 core genes accounting for 91% of the average total gene count per strain. Similarly, four species—*M. smegmatis, M. fortuitum, M. chelonae, and M. phocaicum*—also have high proportions of core genes, making up 87%, 67%, 64%, and 64% of their total gene counts, respectively. Conversely, species like *M. marinum* and *M. intracellulare* display a more balanced distribution of core and dispensable genes, with core genes accounting for 50% and 45% of the gene counts, and dispensable genes making up 48% and 52%, respectively. Interestingly, *M. avium, M. kansasii, and M. abscessus* show a significantly lower proportion of core genes, with 19%, 25%, and 43%, respectively, highlighting considerable genomic flexibility in these species. This pan-genome analysis underscores the substantial genetic diversity and evolutionary dynamics of NTM species, providing a valuable reference for ongoing research and clinical applications.

Additionally, the average gene count per strain for each species ranges from 4,500 to 6,500, further highlighting the diversity among these species. *Mycobacterium (M.) avium* and *M. abscessus* have lower average gene counts per strain, with 4,835 and 4,998 genes, respectively, while *M. smegmatis* and *M. fortuitum* have over 6,000 genes per strain, with 6,550 and 6,316 genes, respectively.

### Phylogenetic characteristics of NTM species

Phylogenetic analyses were performed to explore the evolutionary traits of NTM species and the relationships between different subtypes, offering insights into their pathogenicity and transmission. The phylogenetic analysis (**Figure S1**) revealed that NTM species clustered into two major clades: a slow-growing clade containing *M. avium, M. intracellulare, M. marinum, M. smegmatis, M. fortuitum, M. ulcerans*, and *M. kansasii* with shorter branch lengths, and a rapid-growing clade comprising *M. abscessus and M. chelonae* with longer branch lengths. Here, we primarily focused on the three most prevalent species: *M. abscessus, M. avium, and M. intracellulare* **(Figure 5)**.

**Figure 5.**
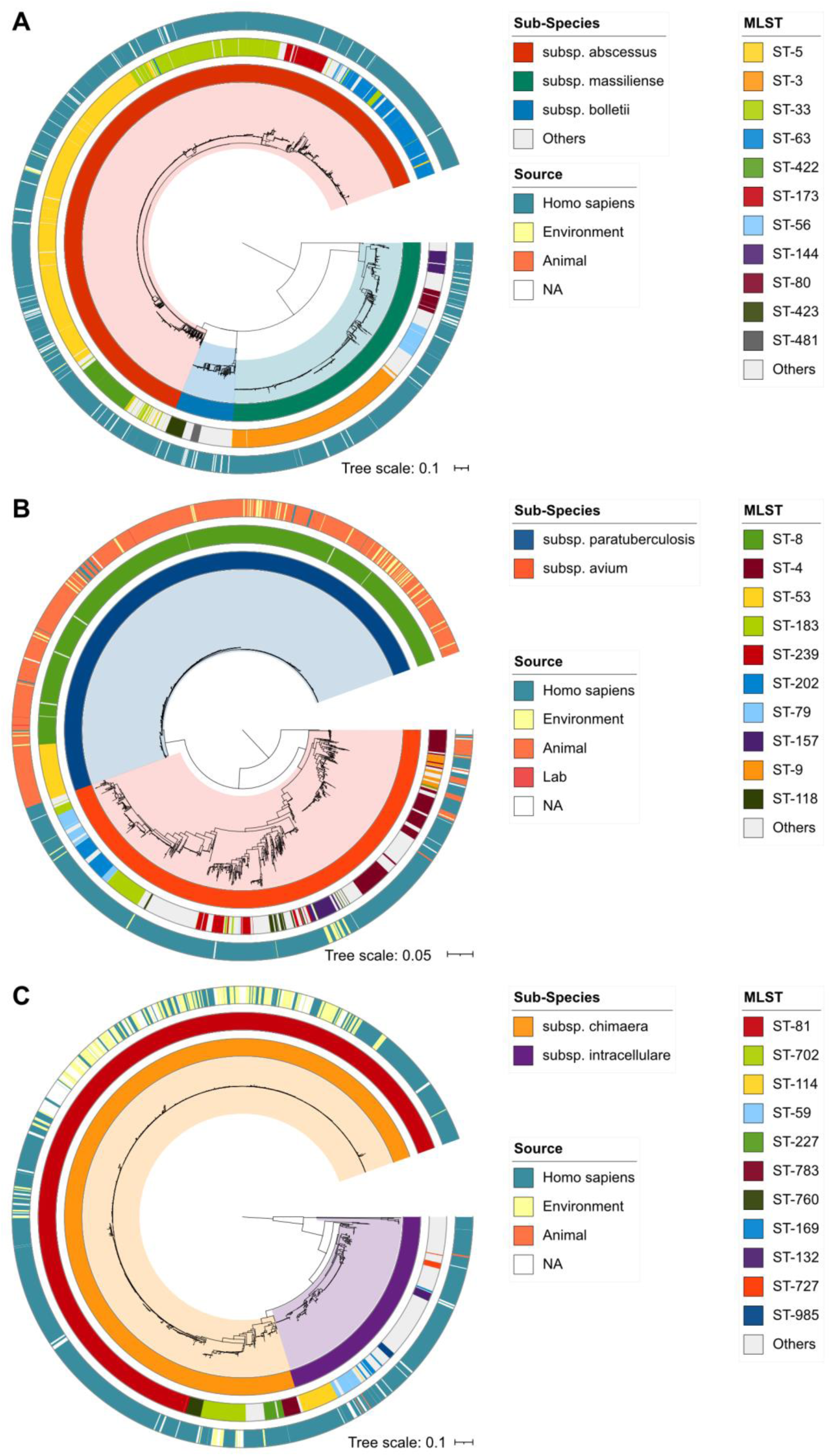
Phylogenetic tree of the three most prevalent NTM species. The evolutionary relationships are depicted for **A.** *Mycobacterium abscessus*, **B.** *Mycobacterium avium*, and **C.** *Mycobacterium intracellulare*, illustrating the main subspecies and their corresponding sequence types (STs).

M. abscessus has evolved into three main clades, corresponding to its subspecies: M. abscessus subsp. abscessus, M. abscessus subsp. massiliense, and M. abscessus subsp. bolletii. M. abscessus subsp. abscessus is the most common, with prevalent ST types ST-5 (49.4%) and ST-33 (25.3%). M. abscessus subsp. massiliense is dominated by ST-3 (70%), while M. abscessus subsp. bolletii primarily features ST-423 (32.3%) and ST-481 (14.5%). All three subspecies are predominantly isolated from human sources, indicating their pathogenicity in humans.

*M. avium* consists of two subspecies: *M. avium subsp. paratuberculosis* and *M. avium subsp. avium*. *M. avium subsp. paratuberculosis* is highly conserved, with two main ST types, ST-8 (91.2%) and ST-53 (8.8%). In contrast, *M. avium subsp. avium* shows greater evolutionary diversity, with ST-4 (17.9%) and ST-183 (8.9%) being the most common. *M. avium subsp. paratuberculosis* primarily infects ruminants, with most strains isolated from animals, while *M. avium subsp. avium* is mainly isolated from humans.

M. intracellulare is divided into two subspecies: M. intracellulare subsp. intracellulare and M. intracellulare subsp. chimaera. M. intracellulare subsp. intracellulare appeared earlier and showed more differentiation, while M. intracellulare subsp. chimaera is more conserved, with ST-81 (87.4%) as the dominant type. These subspecies are primarily isolated from humans, although many M. intracellulare subsp. chimaera strains are found in patient environments.

These analyses offer an enhanced understanding of the evolutionary pathways and pathogenic characteristics of NTM species, serving as an important reference for further research and clinical applications of NTMs.

### Drug resistance profiles of NTM

Our analysis identified resistance genes in 123 species, related to 28 different drugs **(Figure 6A)**. The penam and macrolide resistance-associated gene *mtrA* were the most common, found in 121 species (9,592 strains). This was followed by rifampicin resistance-associated genes, *RbpA,* and *ropB2*, detected in 123 and 120 species, respectively (9,583 and 9,571 strains respectively). On average, each species carried about five resistance genes. Notably, *M. abscessus* had the highest number of resistance genes, totalling 71, with *M. smegmatis* and *M. peregrinum* following with 13 and 12 resistance genes respectively.

**Figure 6.**
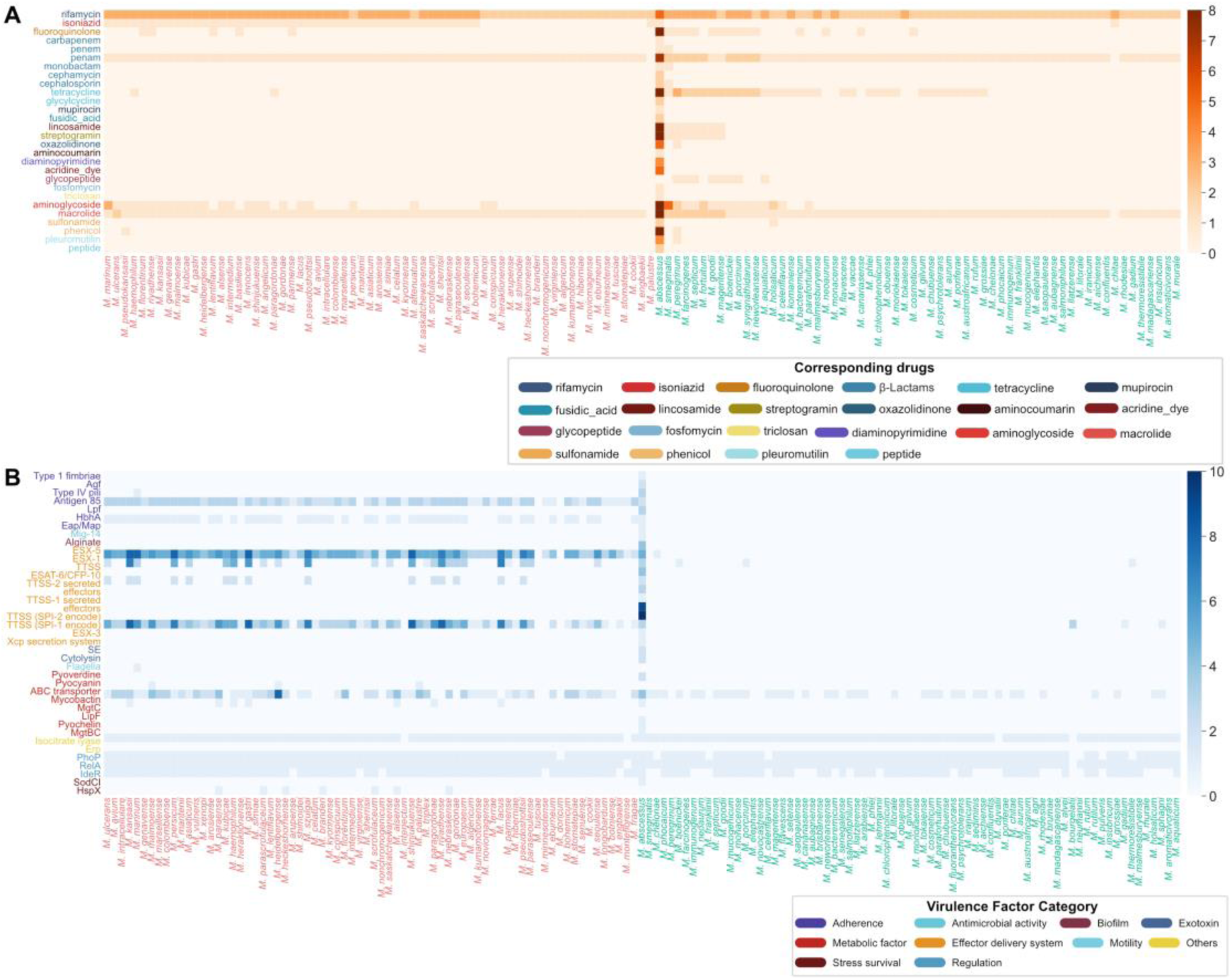
Virulence and Drug Resistance Gene Features in NTM Species. This figure illustrates the distribution of virulence and drug-resistance genes across NTM species, differentiated by growth characteristics. Slowly growing mycobacteria (SGMs) are marked in red, and rapidly growing mycobacteria (RGMs) in green. **A**. Shows the count of resistance genes across species, categorized by the drugs they confer resistance to. **B.** Displays the count of virulence genes across species, categorized by virulence factor categories.

Further comparisons between rapidly growing *mycobacteria* (RGMs) and slowly grown *mycobacteria* (SGMs) revealed distinct patterns in resistance gene distribution. RGMs generally had a broader spectrum of resistance genes, averaging six resistance genes per species, compared to about four genes per SGM species. Although SGMs typically possessed fewer resistance genes, resistance-associated genes for isoniazid (a commonly used antibiotic in tuberculosis treatment) are observed in 45 SGM species, while only two RGM species, *M. abscessus* and *M. chitae*, showed such resistance.

### Virulence characteristics of NTM

Our analysis identified virulence factor (VF) genes in 156 species, comprising 38 different virulence factors categorized into 9 classes based on their mechanisms and functions **(Figure 6B)**. Among these, regulatory virulence factors such as *RelA*, *PhoP*, *IdeR*, and Isocitrate lyase were the most widely distributed in NTMs, with 154, 148, 124, and 152 species harbouring these genes, respectively. Mycobactin exhibited the highest diversity with 14 different virulence genes and was also prevalent in 101 species. It was followed by the effector delivery system category, which includes virulence factors ESX-1, ESX-3, and ESX-5, featuring 12, 10, and 10 virulence genes, respectively.

A comparative analysis between rapidly growing mycobacteria (RGMs) and slowly grown mycobacteria (SGMs) has unveiled distinct trends. SGMs typically display enhanced virulence with a broader spectrum of virulence factors (∼29.3 per species), while RGMs generally exhibit a more limited array (∼4 per species). Notably, effector delivery system-related virulence factors such as ESX-1, ESX-3, ESX-5, ESAT-6/CFP-10, Adherence-related factors Antigen 85, and *HbhA* are widely present in SGMs: ESX-1 in 33 species, ESX-3 in 73, ESX-5 in 78, ESAT-6/CFP-10 in 22, Antigen 85 in 77, and *HbhA* in 56. Conversely, aside from *M. abscessus*, few rapid-growing mycobacteria (RGM) species possess these virulence factors. This may explain why *M. abscessus* exhibits higher pathogenicity. Additionally, the virulence factor Mycobactin shows greater diversity in SGMs, with approximately 14 genes, compared to about 4 in RGMs.

### The web interface of NTM-DB

NTM-DB (https://ngdc.cncb.ac.cn/ntmdb) offers a user-friendly interface, featuring five modules: browse, MLST, search, online analysis, and download.

### Browse module for retrieving comprehensive information for NTM

This database is composed of three main interconnected components: species, strains, and genes. Each component is equipped with primary-level browsing tables, secondary-level detailed information pages, and search functionality **(Figure 7)**, collectively forming a complete framework for exploring NTM.

**Fig. 7.**
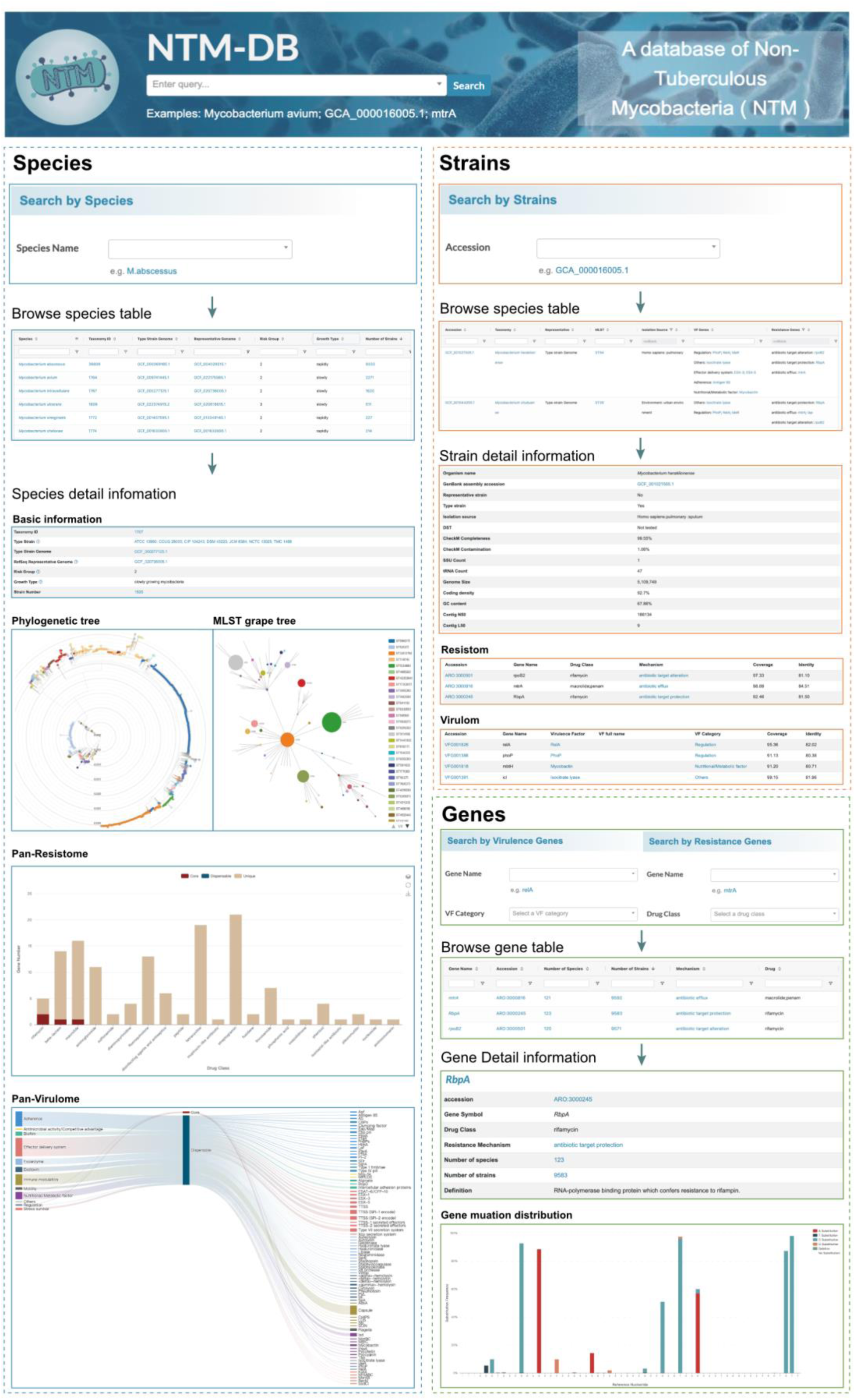
Interface demonstration of NTM-DB for browsing and searching. The top section features a quick search bar for instant access to the homepage. Below, the interface is divided into three main modules: Species, Strains, and Genes. Each module starts with a search interface, followed by detailed data displays and analytical visuals. Arrows indicate the workflow from search queries to the detailed data views within each module.

The species component acts as the database’s foundation, offering a catalogue of NTM species. It provides data on collected strain numbers, taxonomical standard strains, genomic-representative strains, and their corresponding biosafety levels, all displayed in the primary browsing table. Further, the secondary-level page offers detailed species information and interactive visual analyses. These analyses include a whole-genome SNP phylogenetic tree with various display methods and optional metadata colour coding, MLST clustering diagrams that visually represent genetic distances (link length) and strain numbers (node size) to highlight interspecies relationships and diversity, pan-resistance gene bar charts showing the number and conservation of resistance genes across different drug categories, and pan-virulence gene Sankey diagrams detailing gene numbers, conservation levels, and associated virulence mechanisms.

The strain component provides a metadata-enriched interface that lists all matching resistance and virulence genes for each strain, including information on related mechanisms, gene identity, and coverage. This component is vital for researchers focusing on deciphering antibiotic resistance and pathogenicity patterns.

Finally, the gene component catalogues each gene by name, accession number, and prevalence among species and strains in the primary table. The secondary page offers detailed gene information and visualization of gene mutation distribution, providing essential resources for understanding the evolutionary pressures on NTM and their implications for treatment and control strategies.

### User-friendly searching module to access data of interest

NTM-DB offers both quick search and advanced search functions to facilitate access to information on three main database entities: species, strains, and genes, which include both virulence and drug resistance genes. The quick search option on the homepage allows users to find a specific species, strain, or gene rapidly. The advanced search module is organized into four distinct channels: species, strains, virulence factor genes, and drug resistance genes. This structure enables intelligent autofill suggestions and allows for targeted searches by gene category, including searches for virulence factor categories and drug classes associated with resistance genes.

### MLST module

The MLST module in NTM-DB showcases a grape-tree visualization of MLST genotypes for all NTM species **(Figure 8C)**. Each node in the visualization varies in size according to the number of associated strains and is colour-coded to represent different species. The connecting lines between nodes represent the genetic distances, illustrating the relationships among genotypes and species clearly.

**Figure 8.**
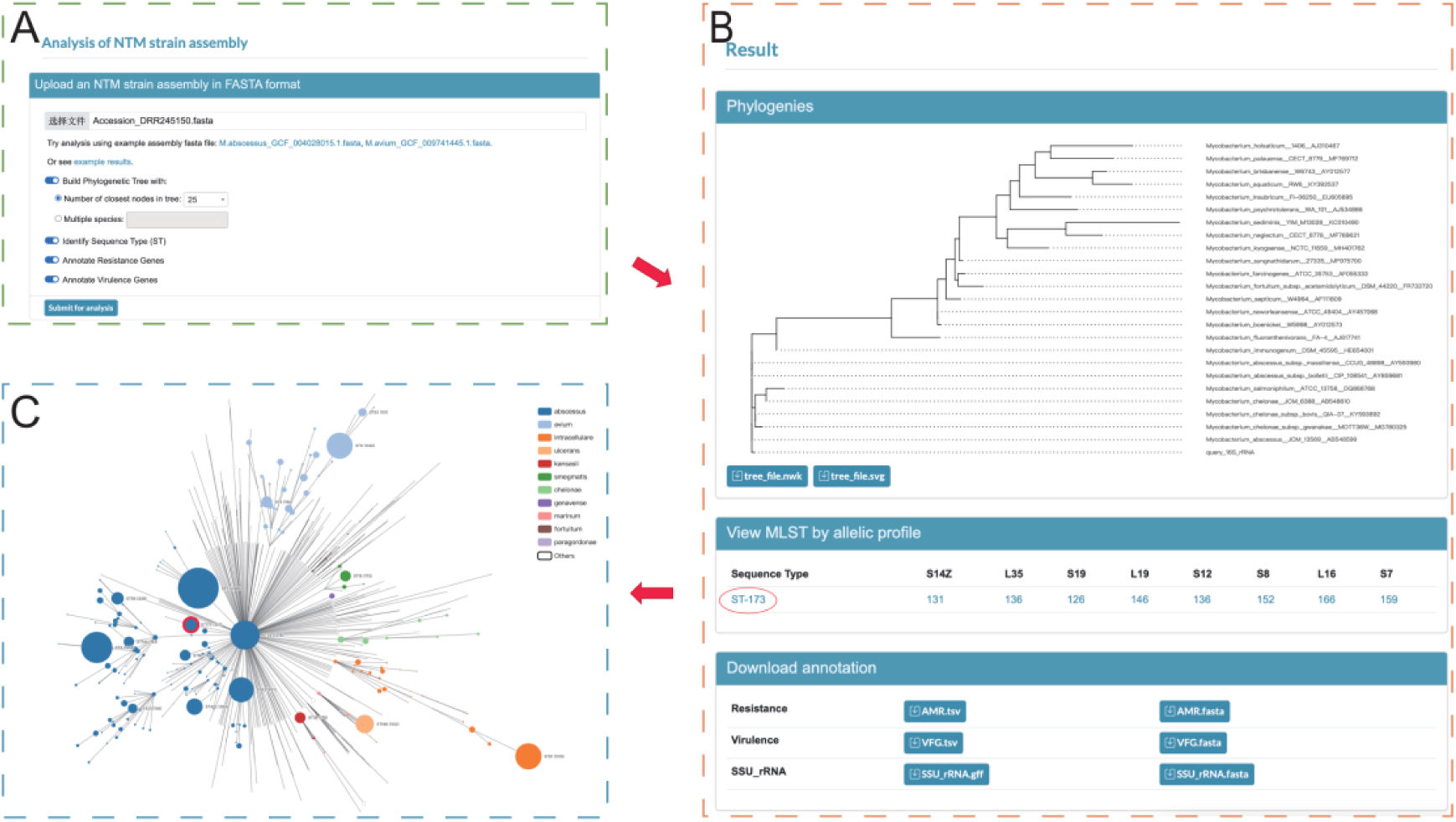
A Case Study of the Analysis Module in NTM-DB. A. The submission interface for uploading NTM strain assemblies in FASTA format. B. The results interface includes detailed outputs such as phylogenetic trees and MLST genotype results. **C.** Visualization of MLST genotypes within the MLST module, with the corresponding sequence types (STs) highlighted in red circles for clear identification.

Interactive features enhance user engagement: hovering over a node displays the strain count while clicking on a node provides access to a detailed table of all corresponding strains. These features not only enhance understanding of intraspecies genetic diversity but also integrate strain MLST annotations from the analysis module, facilitating further exploration of related strains. The utility of this module is further exemplified in a subsequent case study.

### Online analysis module and a case study

The Online Analysis module in NTM-DB significantly enhances its utility by providing advanced features for phylogenetic analysis, MLST genotyping, and resistance and virulence factor gene annotation. This module allows researchers to upload and analyze their NTM strain genome assembly data using a user-friendly interface that integrates various bioinformatic tools for comprehensive analysis.

For convenience and customization, each feature within the module includes a toggle switch, enabling researchers to select specific analyses according to their research needs. Our phylogenetic analysis offers two approaches: researchers can select the number of species for comparison, where the system identifies the genetically closest species via Average Nucleotide Identity (ANI), or they can manually choose specific NTM species based on their research focus. This flexibility allows for the construction of precise phylogenetic trees tailored to different research requirements.

To demonstrate the capabilities of our online analysis tool, we present a case study using the publicly available NTM strain DRR245150 from the SRA database **(Figure 8)**. In this example, the phylogenetic analysis was set to compare the top 25 NTM species. The system calculated the ANI with the type strains of all NTM species and selected the top 25, such as *Mycobacterium (M.) abscessus*, *M. chelonea*, and *M. fortuitum*, for constructing the phylogenetic tree. This tree effectively shows the genetic closeness of the uploaded strain to these species, especially highlighting its close relationship with the *M. abscessus* type strain JCM 13569.

Further MLST genotyping analysis identified DRR245150 as ST-173. A direct link to the MLST module is provided, displaying the node corresponding to ST-173 and facilitating exploration of its relationships and distances to other MLST nodes. Detailed information on the alleles for the eight loci defining ST-173 is also available, with links (*e.g.*https://pubmlst.org/bigsdb?db=pubmlst_mycobacteria_seqdef&page=alleleInfo&locus=S14Z&allele_id=131) to the PubMLST website for comprehensive data on each allele type, facilitating comparisons with other strains sharing the same alleles.

Regarding genetic characteristics, strain DRR245150 contains four resistance genes, including two genes (*RbpA* and *rpoB2*) associated with rifampicin resistance, *mtrA* against macrolides and penams, and *Erm41* against lincosamides, macrolides, and streptogramins. Additionally, four virulence factors were identified in strain SRR245150, including the regulatory genes *phoP* and *relA*, the metabolic factor mycobactin, and the gene *icl* related to isocitrate lyase activity. These annotations are available for download in TSV format and as FASTA sequences in the ’Download Annotation’ section. The SSU rRNA genes used in the phylogenetic analysis are also provided.

It’s important to note that our analysis module prioritizes user privacy, requiring no ancillary personal information and ensuring all data is deleted post-analysis to maintain confidentiality and integrity in handling sensitive genomic data

### Data download module

NTM-DB is dedicated to providing global access to curated NTM research data, supporting ongoing studies into the role of NTM in human health. The platform’s Download module offers detailed, species-level data, including MLST genotypes, annotations of virulence and drug-resistance genes, and pan-genome analyses. This enables comprehensive analysis of the datasets. Additionally, NTM-DB enhances user interaction by including a “Download Table” button on all data tables on the browsing page for immediate data retrieval. These features enable researchers to effectively engage with and contribute to the expanding body of NTM knowledge.

## Discussion and future developments

Recently, the increasing prevalence and severity of non-tuberculous mycobacteria (NTM) infections have drawn significant attention [1, 59, 60]. Although some databases provide partial NTM genomic and bioinformatic data, a dedicated comprehensive platform for browsing, MLST profiling, searching, online analysis, and downloading NTM genomic data, as well as resistance and virulence factors, remains lacking. To address this, we present NTM-DB (https://ngdc.cncb.ac.cn/ntmdb), the first database to systematically compile NTM genomic and bioinformatic data. It features an intuitive, visual, and interactive web interface with online analysis tools suitable for users with varying levels of bioinformatics expertise.

Compared to existing microbial databases, NTM-DB stands out in several key ways: it offers the most comprehensive resource dedicated to NTM species, integrating an extensive collection of strain genomes, bioinformatic data, and associated metadata. This focused approach makes NTM-DB a valuable tool for the research community studying NTM. Additionally, the platform enables in-depth genomic analyses, providing insights into genotyping, phylogenetic relationships, and patterns of resistance and virulence genes. The user-friendly visualizations simplify the interpretation of complex data, enhancing the database’s utility for both basic research and clinical studies.

One of the key features of NTM-DB is its set of online analysis tools, which allow users to upload their own genomic data and perform various analyses in comparison to reference genomes. This flexibility accelerates research by facilitating the rapid acquisition of critical information about NTM strains, from genomic profiling to resistance and virulence characteristics.

As part of the National Genomic Data Center (NGDC) [61], NTM-DB adheres to rigorous standards of data management and reproducibility, ensuring the database remains reliable and up-to-date. Future versions of NTM-DB will incorporate emerging multi-omics data, including transcriptomics, proteomics and metagenomics [62–68], which promise to offer deeper insights into NTM biology. Moreover, the growing need for precise molecular diagnostics in clinical settings underscores the importance of advancing tools for pathogen identification and resistance prediction. NTM-DB will continue to integrate cutting-edge diagnostic standards and tools, supporting public health efforts and clinical applications.

In conclusion, NTM-DB is positioned to significantly advance our understanding of non-tuberculous mycobacteria. With its comprehensive data, intuitive interface, and powerful analysis tools, the database will play a crucial role in facilitating research and innovation in the field, supporting both scientific discovery and clinical practice.

## Code Availability

The bioinformatics tools used by the dataset are described under the Methods section. The code used for data analysis is available on GitHub https://github.com/TianyiLu9494/NTM-DB.

## Authors’ contributions

T.L. and C.L. conceived the study and developed the methodology. T.L., C.L., and Y.Z. conducted the investigation. T.L. collected and curated the data, constructed the database, prepared the figures, and wrote the original manuscript. T.L. and H.W. performed the data analysis and developed the website. F.C., C.L. and Y.Z. revised the original manuscript. C.L., Y.Z., H.W., Z.F., X.J., J.W., P.W., Y.H., H.Y., and J.X. validated the website. F.C. and J.X. supervised the project.

## Acknowledgments

This work is supported by the National Key R&D Program of China (No. 2023YFC2604400), National Natural Science Foundation of China (No. 32370656, No. 32300553), Funds for International Cooperation and Exchange of the National Natural Science Foundation of China (Grant No. 32061143024)

## Supplementary materials

**Figure S1.**
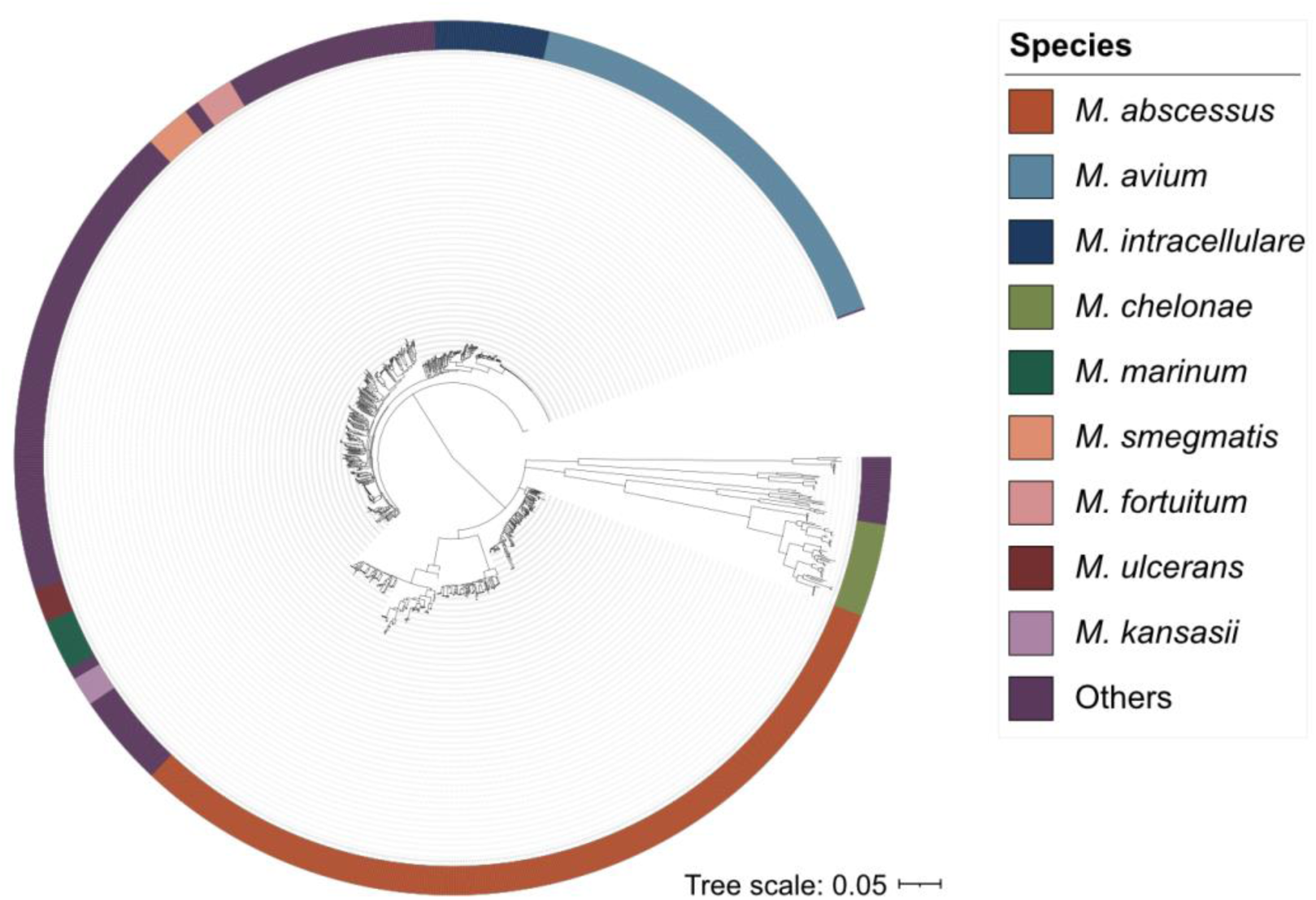
Phylogenetic tree of all NTM species. Comprehensive phylogenetic tree of all identified NTM species constructed using whole-genome SNPs. To reduce redundancy, top 15 highest quality strains per MLST sequence type were retained for abundant species. The nine most prevalent species (*M. abscessus, M. avium, M. intracellulare, M. chelonae, M. marinum, M. smegmatis, M. fortuitum, M. ulcerans, M. kansasii*) are color-coded, while less abundant species are grouped as “Others”.

**File S1 Comprehensive metadata of all NTM strains included in NTM-DB**

Complete metadata information for all collected strains including data source DOI and PubMed ID for traceability and reproducibility.

